## Supplementary Material for "Fiora: Local neighborhood-based prediction of compound mass spectra from single fragmentation events"

### SUMMARY

This Supplementary Material provides additional information on the FIORA model and additional figures and statistics on FIORA's performance. We discuss the impact of collision energies on prediction performance and provide an overview of the dataset with particular focus on the 2022 CASMI challenge. Furthermore, we present example predictions and figures providing a broader overview on similarity scores and biases, and compound class representations of *lipids and lipid-like molecules*.

### ADDITIONAL CHAPTERS

#### The impact of collision energy

The collision energy has a profound impact on the probability of bond breaks and, consequently, fragment ion intensities. FIORA explicitly models collision energies as continuous input values, which are integrated into the fragment ion prediction alongside other covariates after the graph convolution layers. In contrast, CFM-ID predicts spectra at three specific energy levels (10, 20, and 40 eV). ICEBERG does not consider collision energies and, as a result, predicts spectra that represent an average for each compound.

Figure 3 shows the performance of the algorithms at different intervals of collision energies. FIORA's performance declines as the collision energy increases. We suspect that higher collision energies lead to an abundance of higher-order fragments (arising from multiple bond cleavages), which are only partially covered by FIORA's fragmentation algorithm. This assumption is supported by a similar decline in FIORA's peak intensity coverage, shown in Figure 4, at higher collision energy levels. However, ICEBERG exhibits a similar trend to FIORA, with overall lower scores and a smaller decline with increasing collision energy. CFM-ID displays a more rapid decrease in prediction performance, where 10 and 20 eV spectra are predicted with high quality and collision energy levels of 40 eV and above showing significantly worse performance. This indicates that even with multi-step fragmentation, spectra become increasingly harder to predict at higher collision energies. High collision energy spectra are also less prevalent in the training set.

Figure 5 depicts the prediction quality of FIORA for CASMI 16 at different normalized collision energy (NCE) levels. Note that the merged spectrum reflects the experimental setup more closely by modeling the stepped collision energy used in the CASMI 16 challenge, and achieves slightly higher average cosine similarity. Figure 6 shows the prediction quality for CASMI 22 split according to NCE. Here, FIORA predicts spectra significantly better for the low collision energy settings. This is discussed in detail in the Data analysis for CASMI 22 section.

#### Data analysis for CASMI 22

Cosine scores of predicted MS/MS spectra are significantly lower for all algorithms on the CASMI 22 dataset. As noted in the main manuscript, the results must be considered with caution due to inconsistencies between the CASMI 22 data and spectra recorded in the spectral libraries from NIST and MSDial. With that said, we specifically utilized the CASMI 22 dataset to investigate differences in the MS/MS data and to examine the limits of the fragmentation algorithms.

Figure 7 provides an overview of the MS/MS data in the three test sets. The CASMI 22 spectra were obtained at higher collision energies with a significant portion using 50 eV or higher. Such spectra are barely present in the training data. Note that the 10% test split gives a good overview of the distributions in the training set. As all algorithms exhibit worse performance for high collision energies, this explains the low overall cosine scores for CASMI 22 to some degree. FIORA is also more affected by higher collision energies, which explains why this is indeed its worst performing dataset. This can be explicitly seen in Figure 6. At 35 NCE, FIORA predicts spectra at a median cosine similarity of 0.4, which is comparable to that of CFM-ID and ICEBERG. Only at higher NCE values does FIORA's performance decline in comparison. CFM-ID, for example, demonstrates relatively consistent performance across all energy levels.

We also observe a high number of peaks and an abundance of low m/z peaks in the CASMI 22 dataset. In Figure 7, the test split and CASMI 16 have a maximum peak abundance at around 125 m/z, with a steep drop-off for smaller fragments. In contrast, peak abundance for CASMI 22 reaches its maximum for the smallest recorded peaks at around 50 m/z. Furthermore, peak intensity is split among many peaks for CASMI 22. The right column of Figure 7 depicts the number of peaks that account for 80 % of peak intensities. In the case of CASMI 22, intensities are spread among 20 or more peaks for most spectra, indicating a higher number of significant peaks compared to the other datasets. We believe that the high abundance of low m/z peaks, which may cover a substantial amount of the total peak intensity, make CASMI 22 spectra difficult to predict. Interestingly, negative mode spectra have a smaller number of significant peaks, which aligns better with other datasets. This also correlates with an improved performance of Fiora for negative mode CASMI 22 spectra. Overall, the data suggests a considerable structural difference of CASMI 22 spectra compared to the training, test and CASMI 16 spectra. This may also explain the inconsistency between library and CASMI 22 spectra that we reported before.

Most importantly, we observed that algorithms face a severe out-of-distribution problem on CASMI 22. The models learn from the training data, which has a comparable peak m/z distribution to the test split, to distribute the majority of peak intensities among three to five peaks. Consequently, it is unreasonable to expect them to accurately predict spectra where 80% of peak intensities is distributed among 20 or more peaks. It is possible that some form of data pre-processing and noise removal, particularly with regard to low-intensity peaks, could mitigate some of these issues. Importantly though, it is crucial to conduct a thorough data analysis when utilizing any MS/MS prediction software, and it can be concluded that the CASMI 22 dataset is ill-suited for all three algorithms. Fiora is most effective when used in experimental setups that employ low to moderate collision energies, which result in a small number of significant peaks. Stepped collision energies, as seen in CASMI 16, can also be modeled, as Fiora's implementation takes the value of the collision energy as an input.

### Model specifications

**Table 1:** List of molecular and covariate features (on the left), model hyperparameters (center), and training parameters (on the right). Model specifications describe the final set of hyperparameters that were tuned on the validation set.

| | Features |
|---|---|
| Atom | Element |
| | Num of hydrogens |
| | Ring type |
| Bond | Bond type |
| | Ring type |
| Covariates | Collision energy |
| | Molecular weight |
| | Ionization |
| | Instrument type |

| Hyperparameter | Value |
|---|---|
| Graph network type | RGCN |
| Graph layers | 6 |
| Dense layers | 2 |
| Activation function | ELU |
| Embedding dimension | 300 |
| Hidden dimension | 300 |
| Input dropout | 0.2 |
| Latent dropout | 0.1 |

| Training parameter | Value |
|---|---|
| Num epochs | 200 |
| Batch size | 256 |
| Learning rate (LR) | 0.004 |
| Scheduler | Reduce LR on Plateau (patience = 8; factor = 0.5) |

### FIGURES

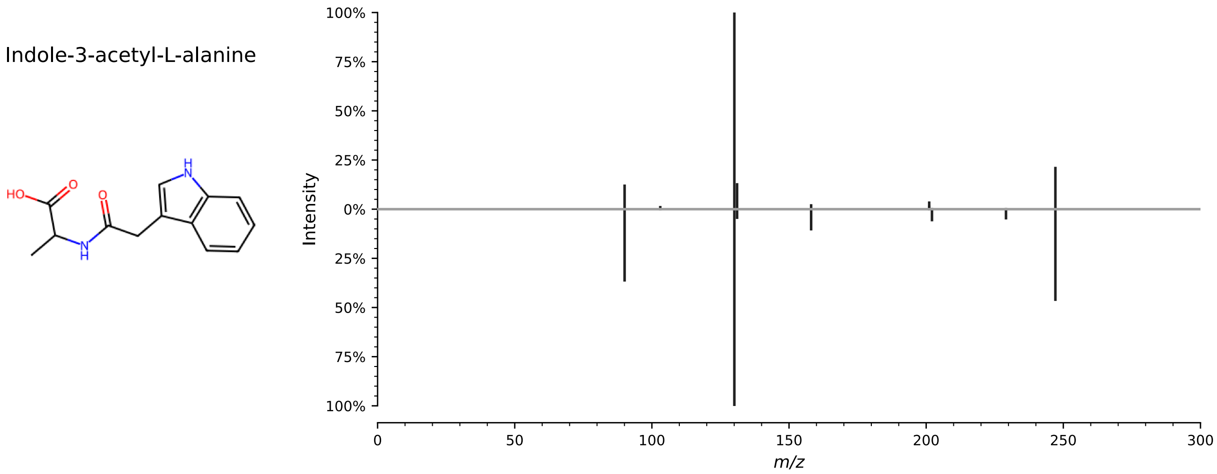

**Figure 1:** Spectral mirror plot for *Indole-3-acetyl-L-alanine* in **positive [M+H]$^+$** ionization mode. This compound is used to illustrate FIORA's fragmentation algorithm in the main manuscript. The upper panel displays the experimental spectrum from the MS-Dial library, while the lower panel shows FIORA's prediction. The cosine similarity between the two spectra is 0.92, and the maximum Tanimoto similarity to the training compounds is 0.73 (using a 2048-bit Morgan fingerprint with a radius of 3).

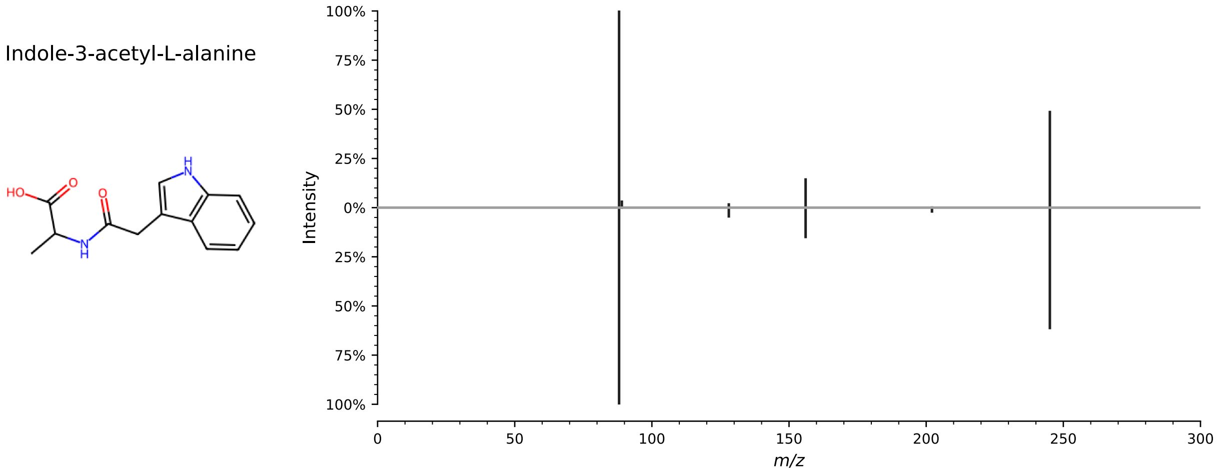

**Figure 2:** Spectral mirror plot for *Indole-3-acetyl-L-alanine* in **negative [M-H]$^-$** ionization mode. This compound is used to illustrate FIORA's fragmentation algorithm in the main manuscript. The upper panel displays the experimental spectrum from the MS-Dial library, while the lower panel shows FIORA's prediction. The cosine similarity between the two spectra is 0.98, and the maximum Tanimoto similarity to the training compounds is 0.73 (using a 2048-bit Morgan fingerprint with a radius of 3).

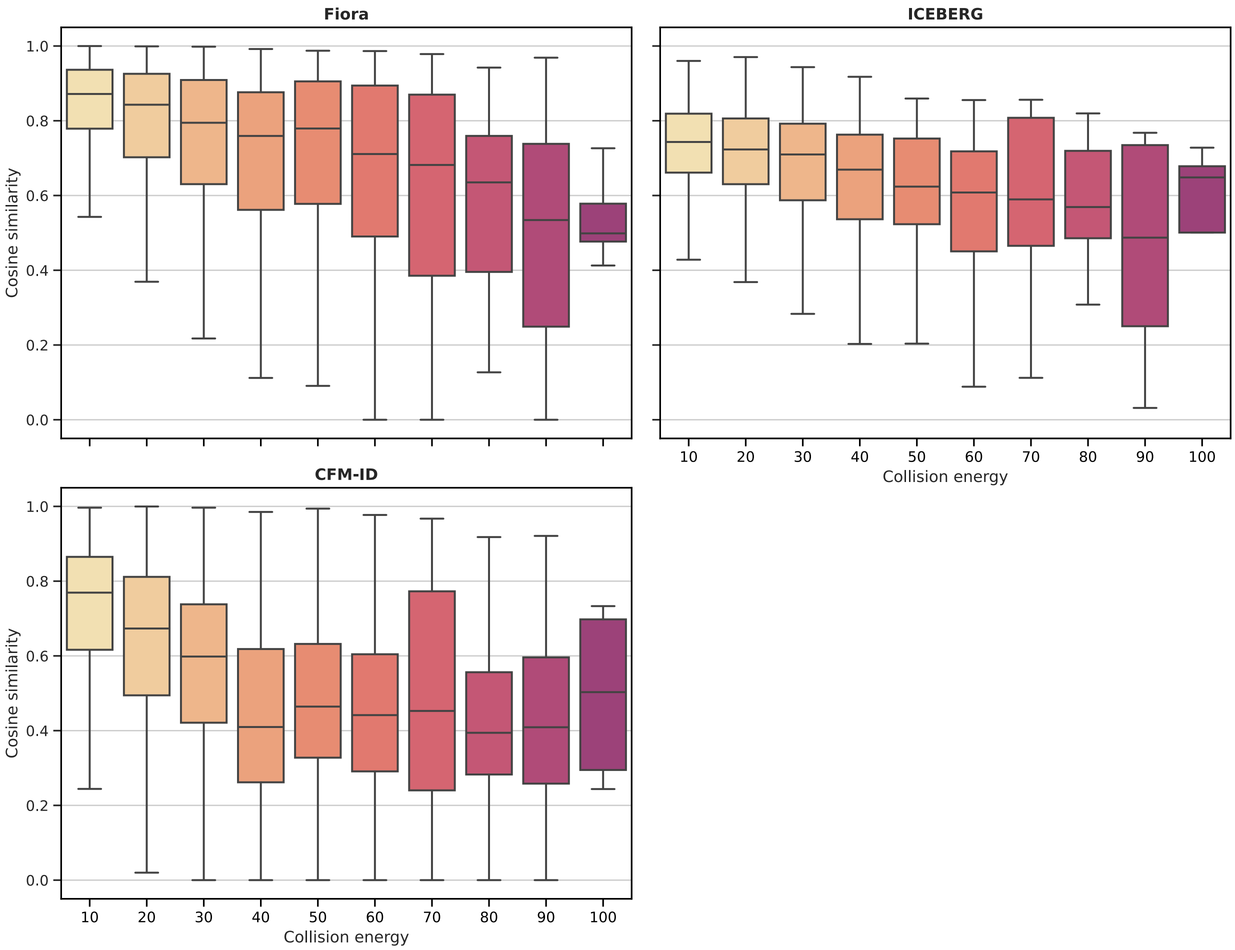

**Figure 3:** Cosine similarity distribution at collision energy intervals. For all algorithms, a decline in prediction performance with increasing collision energy can be observed.

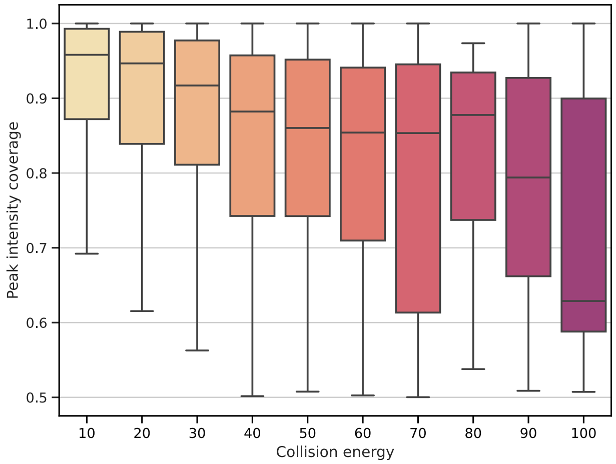

**Figure 4:** Peak intensity coverage at collision energy intervals. Coverage declines with increasing collision energy.

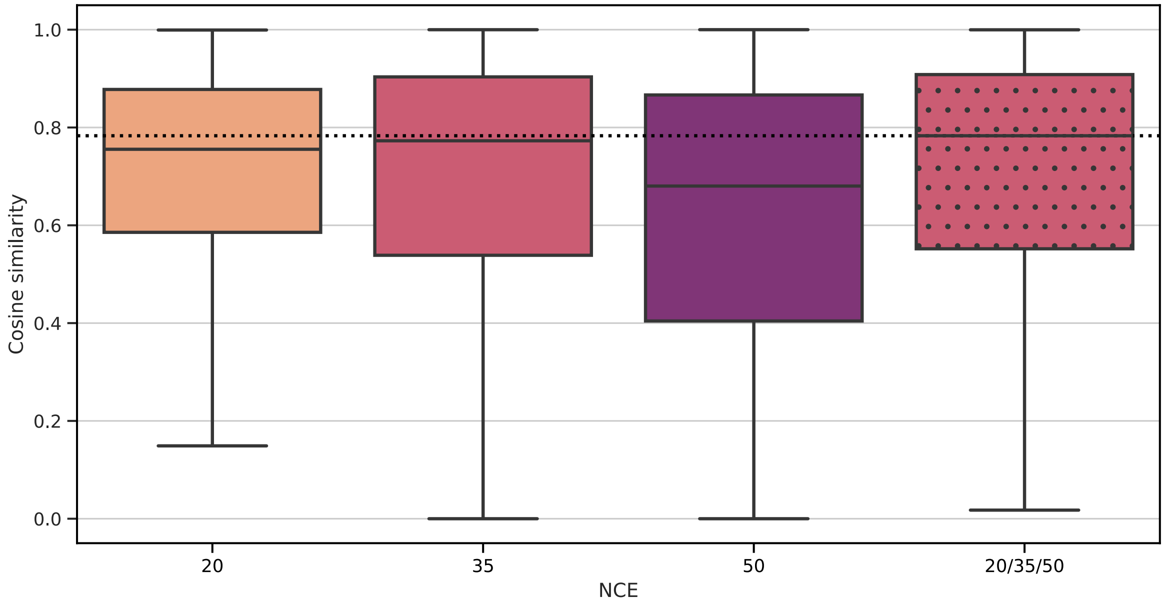

**Figure 5:** Cosine similarity between the CASMI 16 challenge spectra and the predictions at collision energy levels 20, 35, 50 (NCE), and the merged prediction (highlighted). The merged spectra, which correspond more closely to the experimental setup, have on average a higher cosine similarity than the predictions at the individual steps (compare medians with the dotted line).

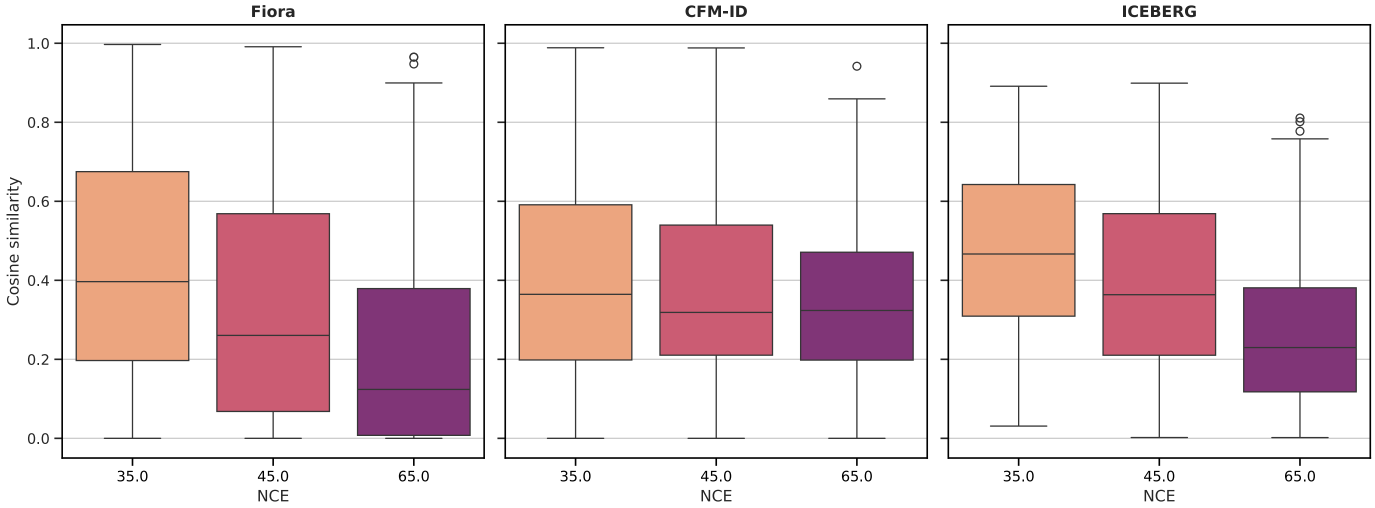

**Figure 6:** A comparison of cosine similarity of CASMI 22 at collision energy levels 35, 45, and 60 (NCE). All algorithms exhibit higher prediction quality at the lower collision energies. This trend is more pronounced for FIORA and barely visible for CFM-ID. At a normalized collision energy of 35, the performance of FIORA is comparable to that of the other algorithms and declines only at the higher energy levels.

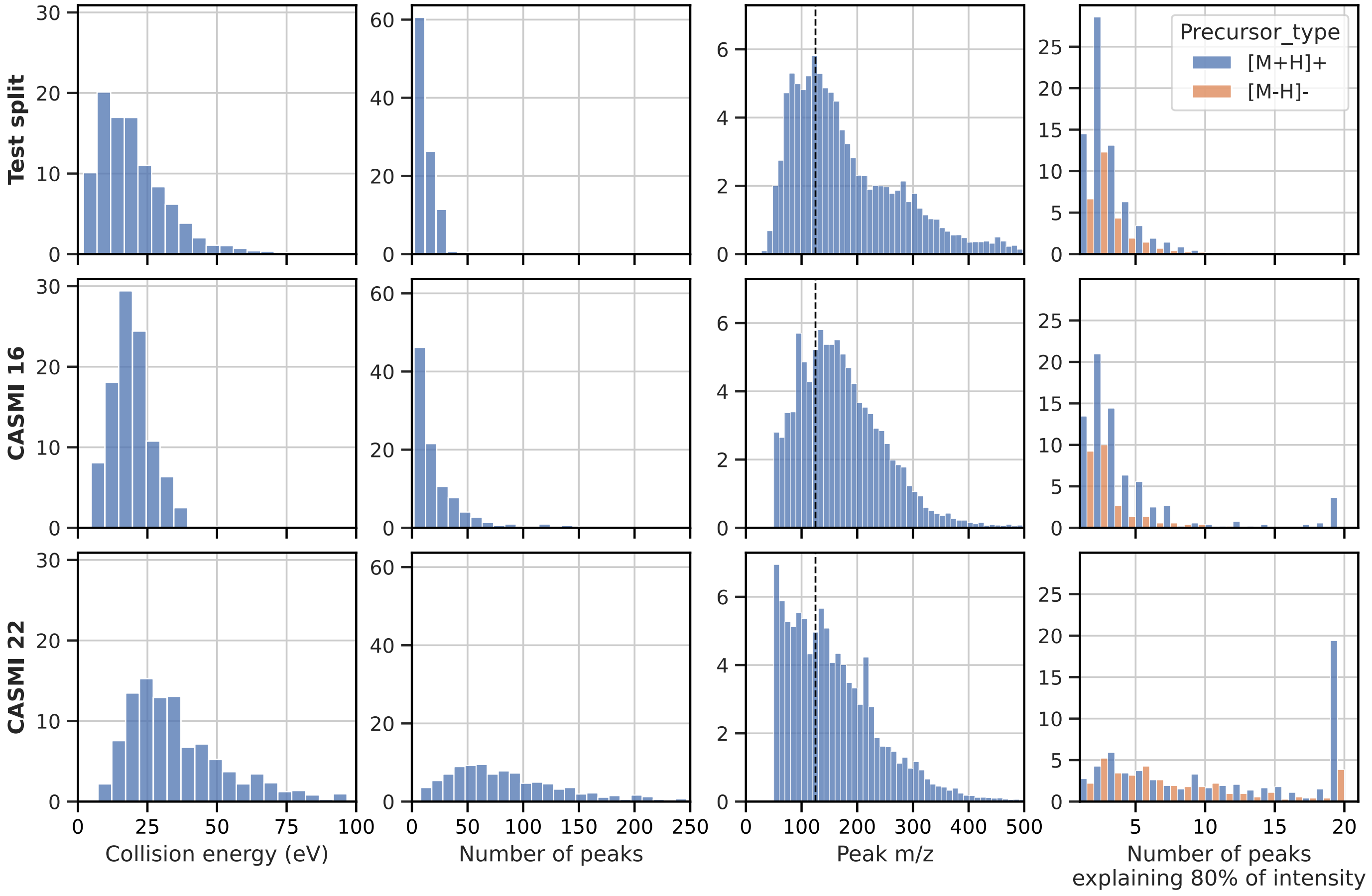

**Figure 7:** Distribution of MS/MS characteristics in the test datasets. The x-axes describe the abundances in percent. The CASMI 22 dataset exhibits distinctively different distributions compared to the Test split and CASMI 16. Collision energies are higher with a larger spread, the spectra contain significantly more peaks, which includes an abundance of low m/z peaks (under 120 m/z). Lastly, peak intensities in CASMI 22 are distributed across a large number peaks, with 80% of peak intensity being explained by 20 or more peaks in many cases.

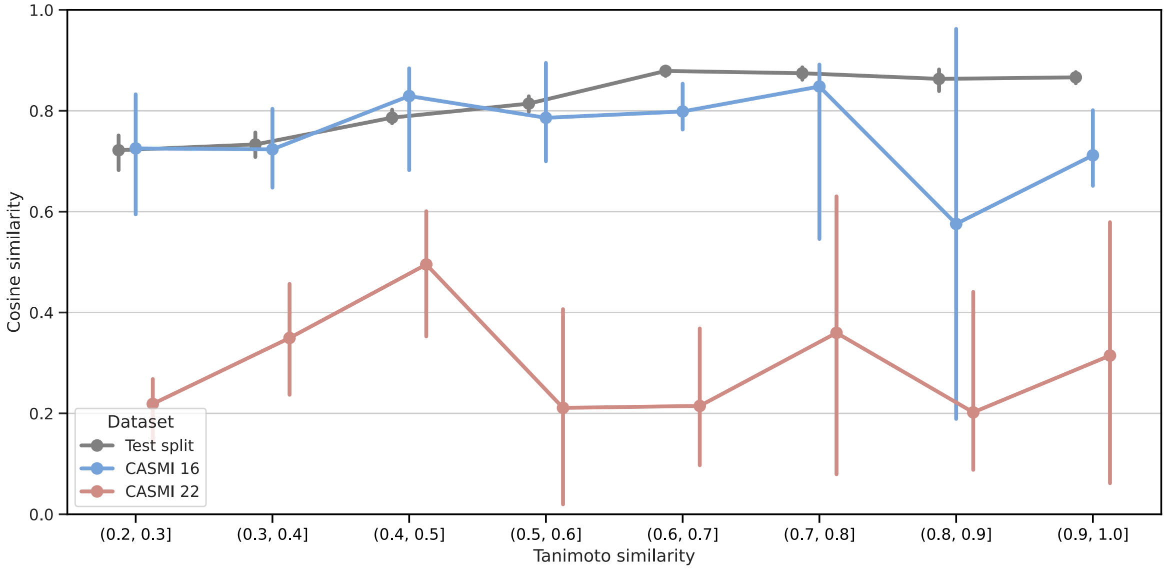

**Figure 8:** Cosine similarity at intervals of structural similarity of compounds from the three test sets to training compounds. Structural similarity was measured by the maximum Tanimoto similarity (Jaccard index) using Morgan fingerprints with 2048 bits and a radius of 3. Spectral predictions were made by FIORA.

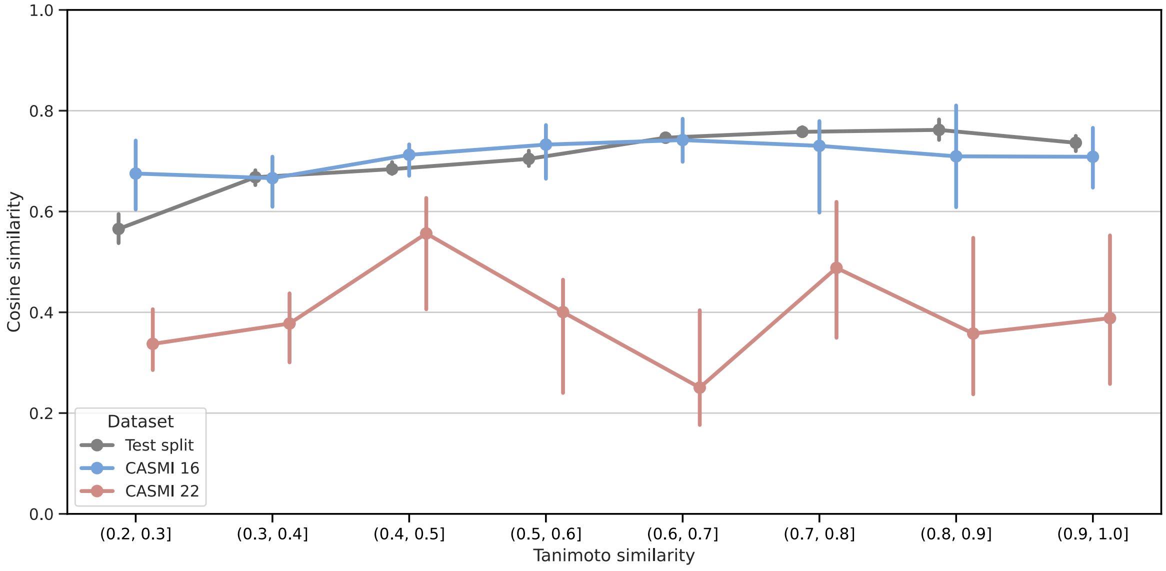

**Figure 9:** Cosine similarity at intervals of structural similarity of compounds from the three test sets to training compounds. Structural similarity was measured by the maximum Tanimoto similarity (Jaccard index) using Morgan fingerprints with 2048 bits and a radius of 3. Spectral predictions were made by ICEBERG.

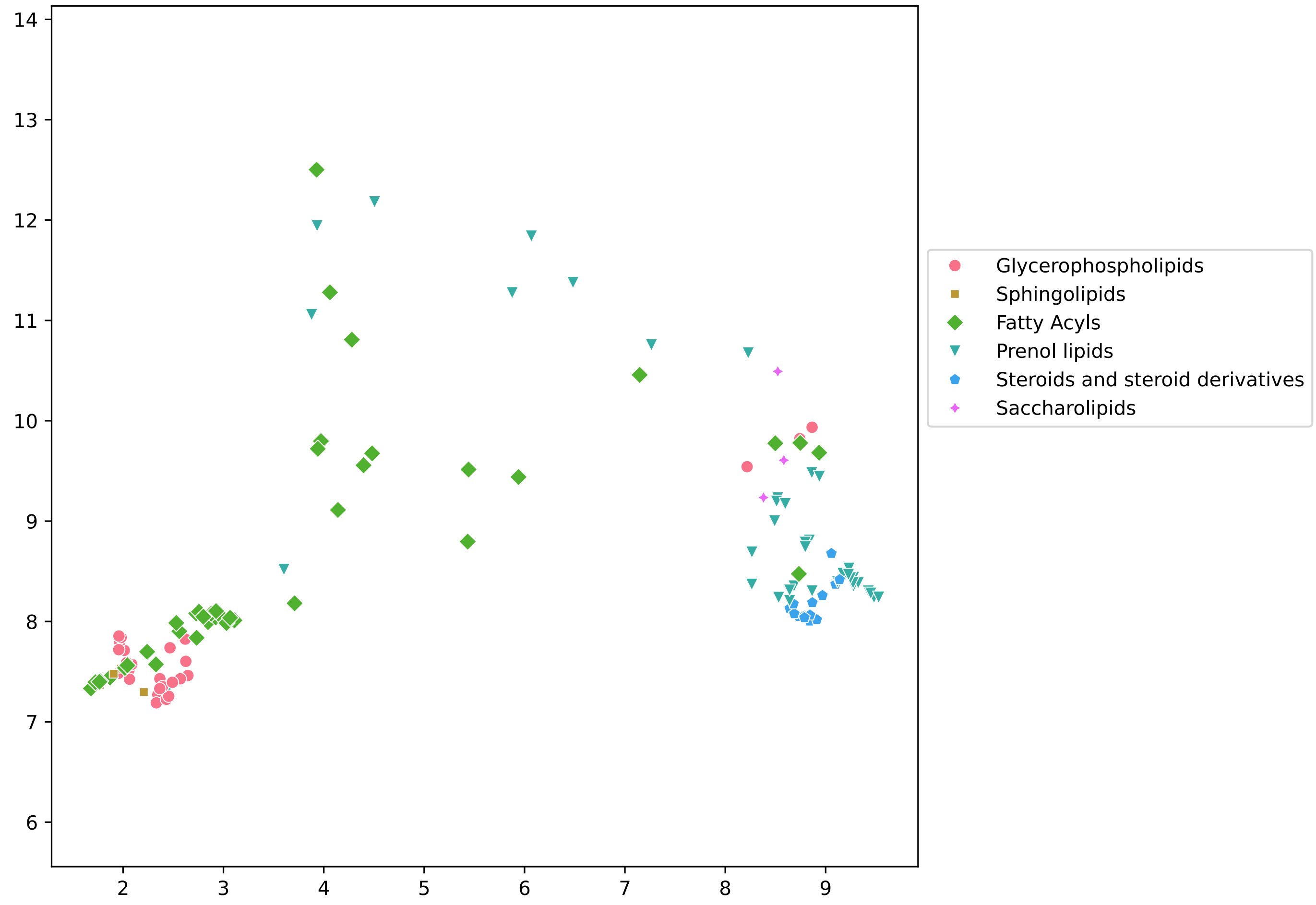

**Figure 10:** UMAP of graph embeddings depicting *lipids and lipid-like molecules* annotated at the compound class level. Each point corresponds to a unique compound and is colored according to compound class, which were annotated by ClassyFire [1]. Dimensionality reduction was performed with respect to all compounds, resulting in a global arrangement of compound embeddings (compared to Figure 11, which depicts a more local representation). Note that the UMAP is not identical to the one presented in the main manuscript, in spite of using the same seed for dimensionality reduction. Compounds from other superclasses were excluded to highlight specifically the arrangement within the *lipids and lipid-like molecules*.

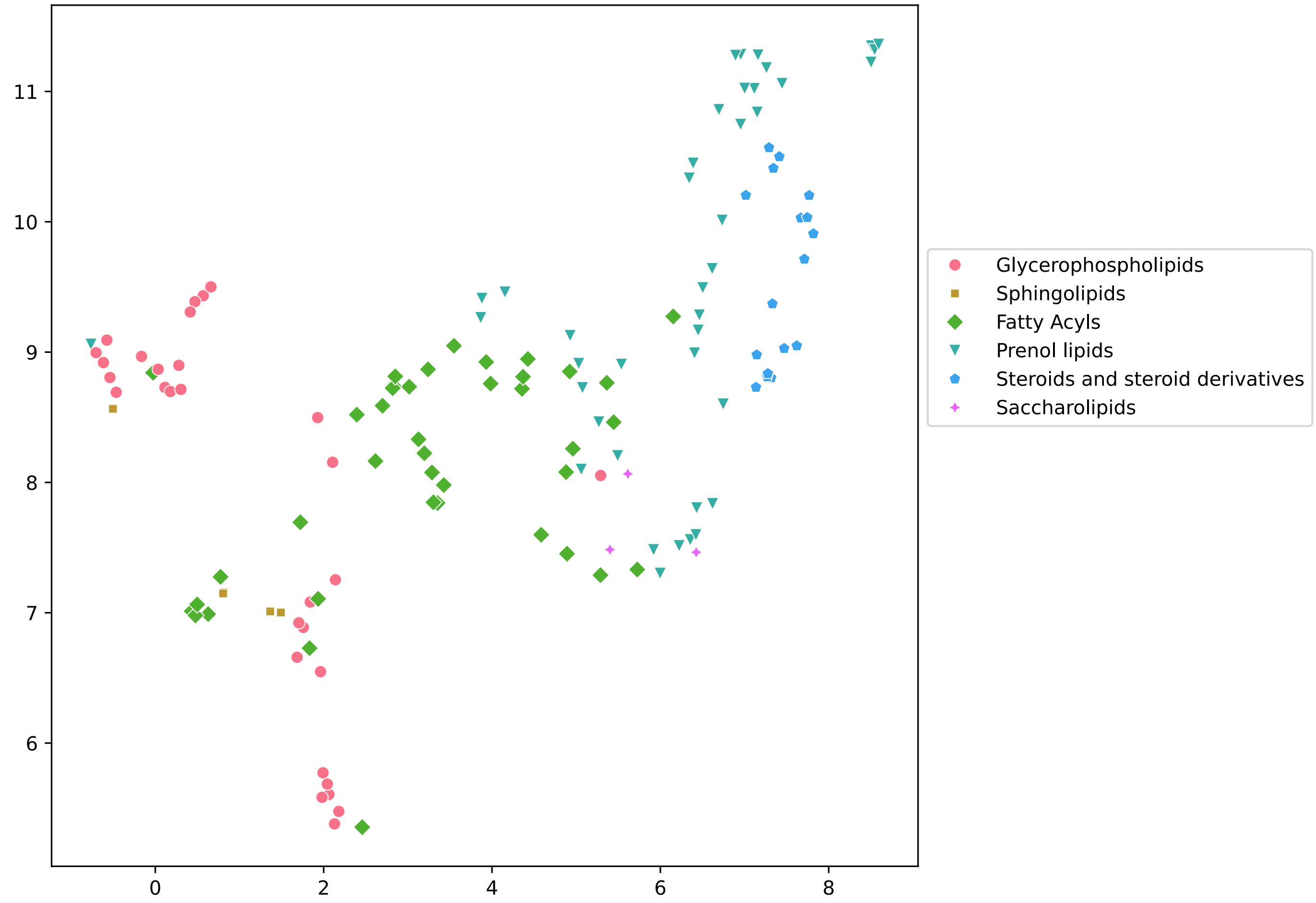

**Figure 11:** UMAP of graph embeddings depicting *lipids and lipid-like molecules* annotated at the compound class level. Each point corresponds to a unique compound and is colored according to compound class, which were annotated by ClassyFire [1]. Dimensionality reduction was performed considering only compound from this superclass. This illustrates a more local arrangement of compound embeddings with respect to other *lipids and lipid-like molecules* (compared to Figure 10, which depicts a more global representation).

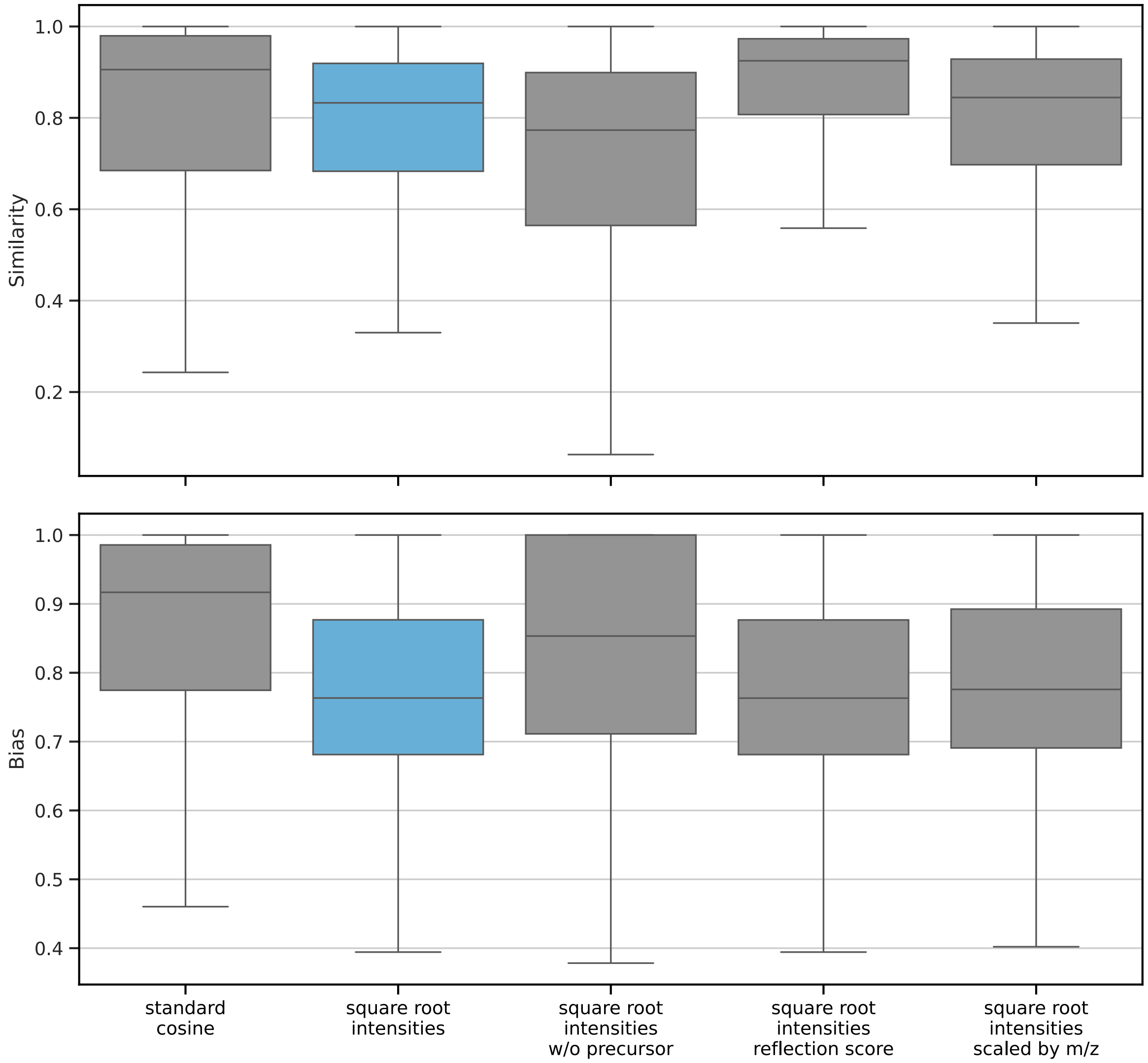

**Figure 12:** Distributions of different similarity scores and their biases evaluated on the test split. Cosine similarity using square root intensities is highlighted because it is the widely preferred spectral similarity score. It also exhibits lower bias values than the standard cosine similarity. As such, it is presumably better at capturing low-intensities peak patterns. Variants of the cosine score include the removal of the precursor peak (w/o precursor), the removal of unmatched (noise) peaks from the query (reflection score), and intensity values scaled by their m/z values. Note that the variants exhibit higher cosine biases (e.g., w/o precursor) or have a narrower range of similarity values (e.g., reflection score). Refer to [2], [3], and [4] for further discussions on spectral similarity scores and biases.
